# Nsp1 stalls DNA Polymerase α at DNA hairpins

**DOI:** 10.1101/2024.09.03.608162

**Authors:** Andrey G. Baranovskiy, Lucia M. Morstadt, Nigar D. Babayeva, Tahir H. Tahirov

## Abstract

The human primosome, a four-subunit complex of DNA primase and DNA polymerase alpha (Polα), plays a critical role in DNA replication by initiating RNA and DNA synthesis on both chromosome strands. A recent study has shown that a major virulence factor in the SARS-CoV-2 infection, Nsp1 (non-structural protein 1), forms a stable complex with Polα but does not affect the primosome activity. Here we show that Nsp1 inhibits DNA synthesis across inverted repeats prone to hairpin formation. Analysis of current structural data revealed the overlapping binding sites for Nsp1 and the winged helix-turn-helix domain of RPA (wHTH) on Polα, indicating a competition between them. Comparison of the inhibitory effect of Nsp1 and wHTH on DNA hairpin bypass by Polα showed an 8-fold lower IC_50_ value for Nsp1 (1 µM). This study provides a valuable insight into the mechanism of inhibition of human DNA replication by Nsp1 during a SARS-CoV-2 infection.

## Introduction

The SARS-CoV-2 coronavirus is the causative agent of the COVID-19 (Coronavirus disease-2019) pandemic, which had a devastating impact on public health and the global economy. SARS-CoV-2 dysregulates the immune inflammatory response and is characterized by organ dysfunction, high level of cytokines, and lymphopenia (1). Nsp1 is a major virulence factor for SARS, which plays an important role in the suppression of the innate immune response (2). The SARS-CoV-2 protein interaction map, published in 2020, shows that the human primosome is a potential target for Nsp1 (3). Intriguingly, among all DNA replication factors, only primosome is targeted by SARS-CoV-2. The cryoEM structure of the complex of human primosome with a small globular domain of Nsp1 revealed that it interacts with an exonuclease domain of Polα (4). Nsp1 is docked away from the active site and the DNA-binding cleft of Polα and showed no effect on its DNA-polymerase activity. Thus, the mechanism of Nsp1 action on human primosome and DNA replication is still unclear.

## Results and Discussion

Recently we have shown that RPA, a single-strand DNA-binding protein, is critical for human primosome during DNA synthesis across inverted repeats and that the flexible wHTH domain of RPA is important for RPA-Polα cooperation (5). We predicted the wHTH-binding site on Polα by the modeling based on high structural similarity between the wHTH and wHTH2 domains of RPA and CST (CTC1/STN1/TEN1), respectively (5). Consistent with that prediction, the AlphaFold-multimer (6,7) generated a very similar model of the wHTH-Polα complex (Fig. 1). Strikingly, the alignment of wHTH/Polα and Nsp1/Polα complexes revealed the overlapping binding sites for wHTH and Nsp1 (Fig. 1). This finding points towards a potential role of Nsp1 in the disruption of the RPA-Polα interaction, which would result in primosome stalling at DNA hairpins.

**Figure 1.**
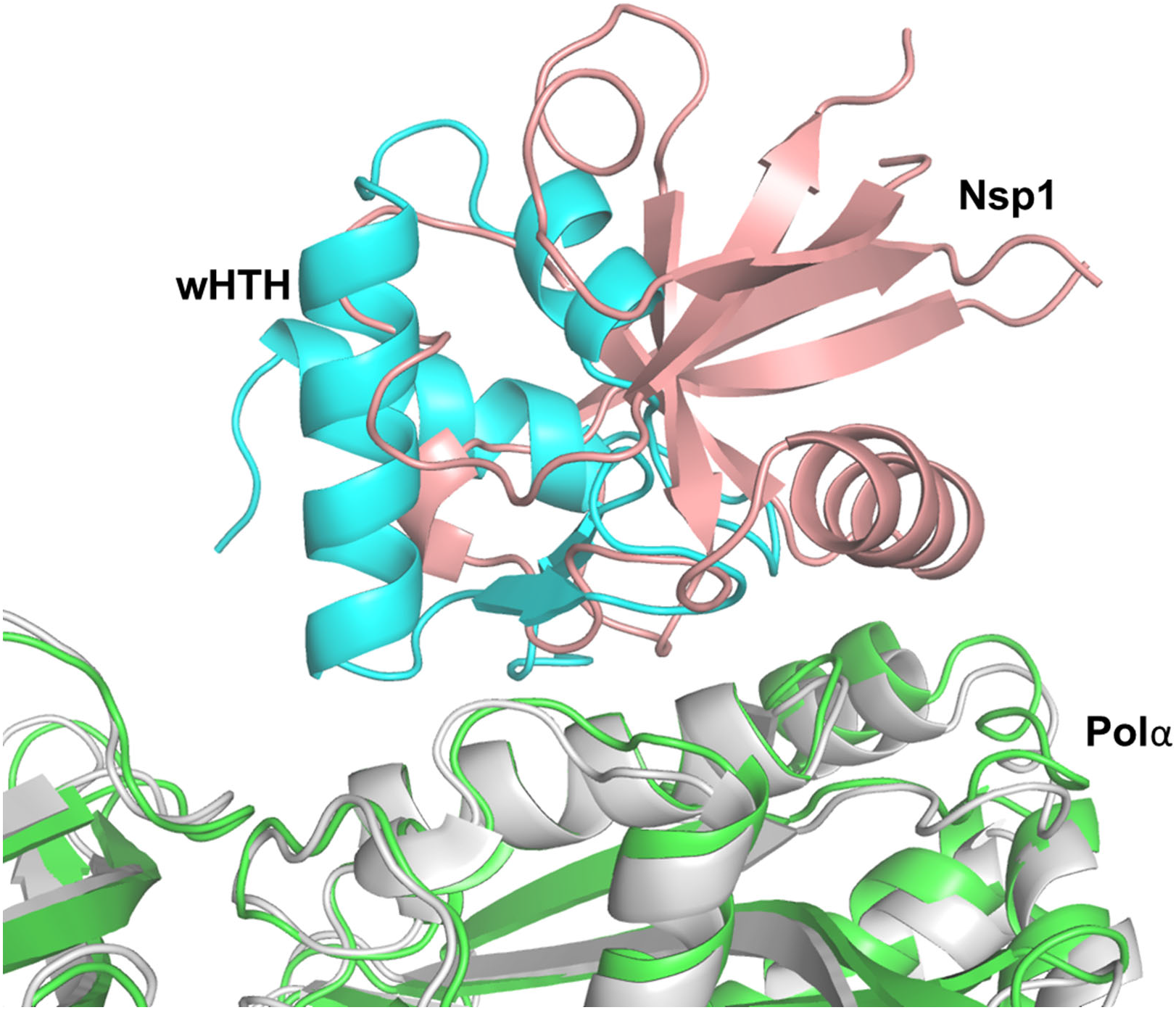
Nsp1 and the wHTH domain of RPA have overlapping docking sites on Pol*α*. The wHTH/Polα (AlphaFold-multimer, ver. 2.0, DeepMind) and Nsp1/Polα (pdb code 7opl) complexes were aligned with rmsd of 1.17 Å using the Polα catalytic domain. Nsp1, wHTH, and Polα in the complex with Nsp1 and wHTH are colored salmon, cyan, gray, and green, respectively. PyMol Molecular Graphics System (version 1.8, Schrödinger, LLC) was used for alignment and figure preparation.

As we showed previously (5), Polα efficiently bypasses the nine base-pair (bp) hairpin in the presence of RPA (Fig. 2, lanes 1-4). The addition of Nsp1 significantly inhibited the hairpin bypass resulting in pronounced Polα pausing at the beginning of a hairpin (Fig. 2, lanes 4-7). The replacement of conserved Val28 by aspartate, a mutation known to disrupt the Nsp1/Polα complex (4), almost completely recovered the hairpin bypass (Fig. 2, lanes 7-9). These results indicate that the inhibitory effect of Nsp1 on DNA synthesis across inverted repeats is mediated by its specific interaction with Polα.

**Figure 2.**
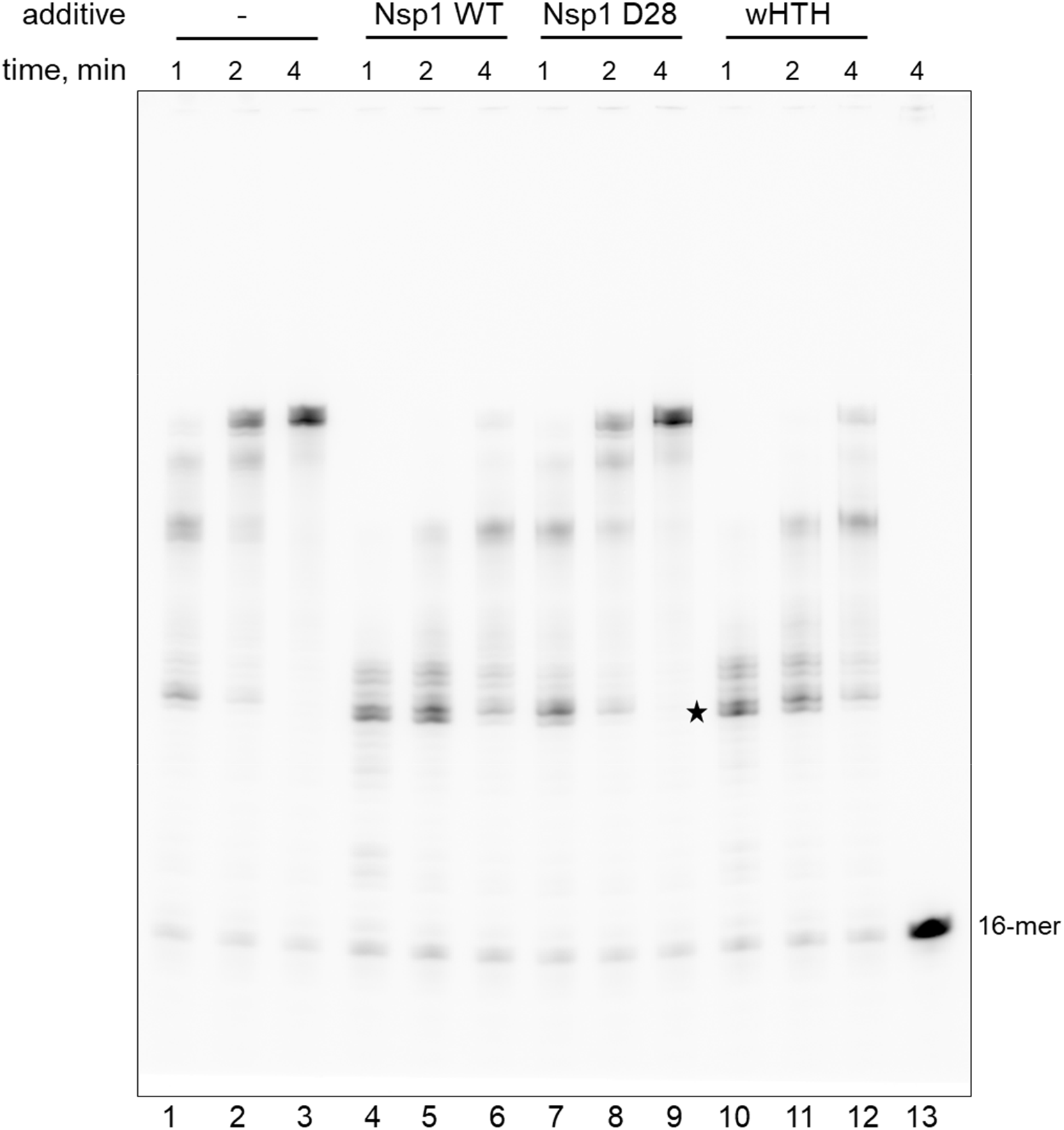
Nsp1 inhibits DNA synthesis across a 9-bp hairpin by the RPA/Pol*α* complex. The DNA-synthetic activity of Polα in extension of a 16-mer DNA primer with a Cy5-fluorophore attached to the 5’-end was tested at 35 °C in a 10 μl reaction containing 0.25 μM template:primer, 20 nM enzyme, 1 µM RPA, 25 μM dNTPs; 8 µM Nsp1 or 40 µM wHTH were added as indicated on the figure. Lane 13 is a control incubation in the absence of proteins. The star denotes the pause site at the beginning of a hairpin located 15 nucleotides from the 3’-end of the primer.

The wHTH domain also inhibits the hairpin bypass but with a weaker efficiency (Fig. 2, lanes 10-12). We compared the inhibitory effect of Nsp1 and wHTH on 9-bp hairpin bypass by the RPA/Polα complex (Fig. 3). The obtained IC_50_ values (1.03 ± 0.04 µM for Nsp1 and 8.14 ± 0.38 µM for wHTH) indicate that *in vivo*, Nsp1 can efficiently compete with the wHTH domain of RPA for interaction with Polα resulting in primosome stalling at hairpins. The effect of Nsp1 on RPA/Polα complex is therefore similar to the deletion of the wHTH domain (5). These findings allow us to speculate that Nsp1 has a strong potential to inhibit synthesis of the telomeric C-strand by the CST/primosome complex because the wHTH2 domain of CST binds Polα at a similar site as Nsp1 (4,8).

**Figure 3.**
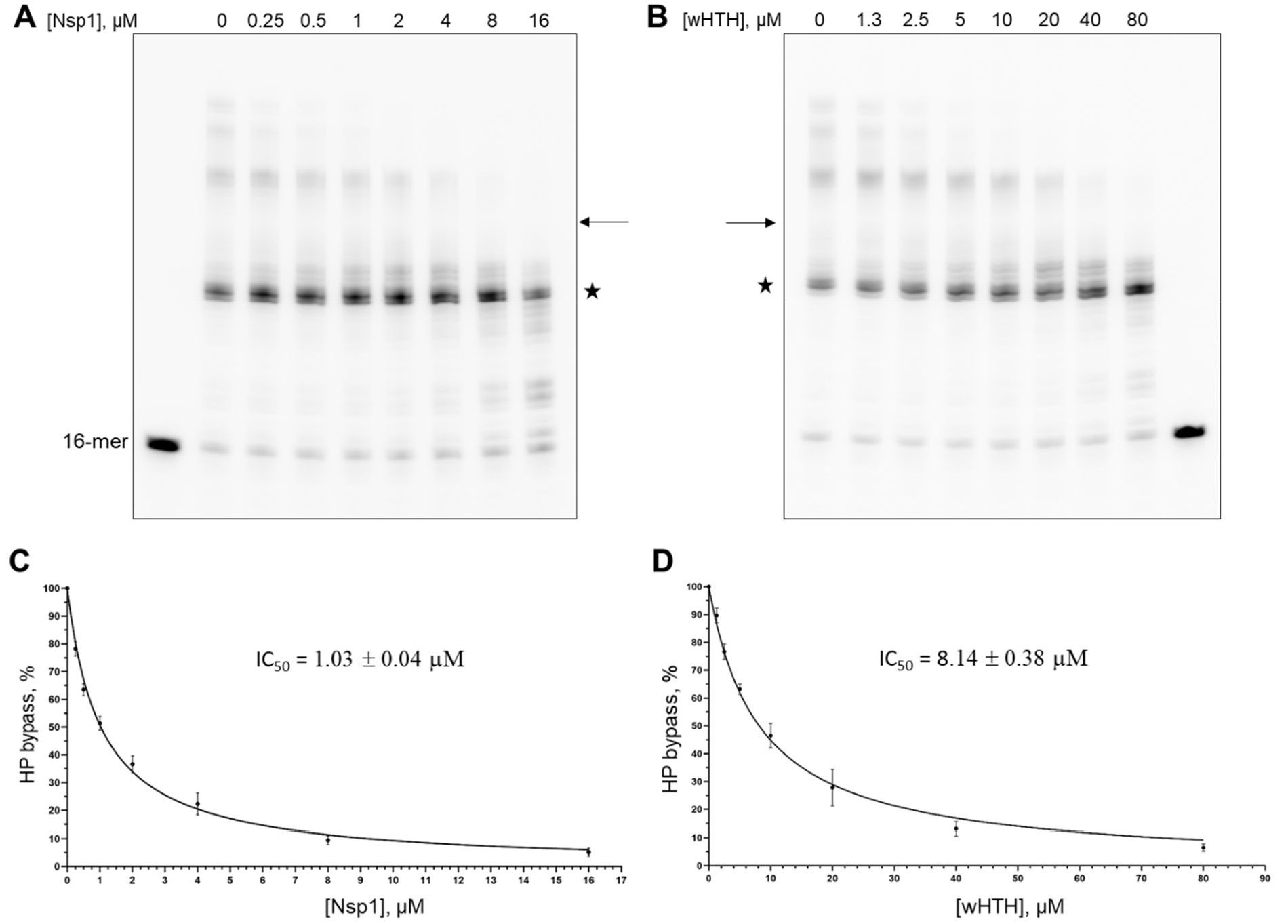
Comparison of the inhibitory effect of Nsp1 and wHTH on hairpin bypass by RPA/Pol*α*. **A** and **B**, representative gels showing titration of DNA-synthetic activity by Nsp1 and wHTH, respectively. The star denotes the pause site at the beginning of a hairpin. The bands above the arrow were counted as products of hairpin bypass. Reactions were incubated for 1 min at 35°C. **C** and **D**, the mean value of the percent of hairpin bypass is plotted against Nsp1 and wHTH concentration, respectively. The error bars denote the standard deviation. The IC_50_ values were calculated from three independent experiments using GraphPad Prism.

A deficiency in the catalytic subunits of primase and Polα causes the X chromosome-linked reticulate pigmentary disorder (XLPRD), which is characterized by the activation of type I interferon signaling, persistent inflammation, and recurrent lung infections (9,10). It was proposed that a reduced level of cytosolic RNA/DNA hybrids triggers this disease, but the molecular mechanism is not clear (10). On the other hand, the inhibited DNA replication may cause damage to genomic DNA and the accumulation of replication fork-derived DNA in the cytoplasm, which then stimulates the production of type I interferons and pro-inflammatory cytokines through the cGAS-STING pathway (11,12). Of note, an abnormal level of cytokines is a hallmark of severe forms of COVID-19, where a dysregulated inflammatory response primarily affects the lungs (1). In addition, by blocking DNA replication, Nsp1 may affect proliferation of T- and B-lymphocytes and, therefore, suppress an adaptive immune response against SARS-CoV-2.

## Materials and Methods

Human RPA and the Polα catalytic domain were obtained as described before (5). The expression vector pMCSG53 encoding for the globular domain of Nsp1 SARS-CoV-2 (residues 13-127) with an N-terminal His-tag followed by the Tev-cleavage site was obtained from Addgene (catalog # 167256) and described in (13). The Nsp1 variant bearing a point mutation V28D was generated by side-directed mutagenesis. The pASHSUL-1 vector for expression of the wHTH domain was made by inserting the coding sequence for the RPA32 residues 178-270 downstream the sequence encoding for the His-Sumo tag (14). Nsp1 and wHTH were expressed in the *Escherichia coli* strain BL21 (DE3) at 16 ºC for 15 hours following induction with 0.05% α-lactose and 0.3 µg/ml anhydrotetracyclin, respectively. Nsp1 and its V28D variant were purified to near-homogeneity (Fig. S1) in two chromatographic steps using Ni-IDA column (Bio-Rad) and Q HiTrap (Cytiva) columns. After the first step, the His-tag was removed by Tev protease during overnight digestion on ice. The wHTH domain of RPA was purified in a similar way, with a His-Sumo tag being digested by dtUD1 protease (14) and removed from the sample by the pass through a HisTrap column (Cytiva). The purified proteins were dialyzed against buffer containing 20 mM Tris-Hepes, pH 7.8, 150 mM NaCl, 2% glycerol, 1 mM TCEP, concentrated to ∼1 mM, and frozen in 10 µl aliquots. Protein concentrations were estimated by measuring the absorbance at 280 nm and using the extinction coefficients calculated with ProtParam (15).

DNA-synthetic activity of Polα was tested in 10 μl reactions containing 0.25 μM template:primer, 20 nM Polα, 1 µM RPA, 25 μM dNTPs, and the buffer: 30 mM Tris-Hepes, pH 7.8, 120 mM KCl, 30 mM NaCl, 1% glycerol, 1.5 mM TCEP, 5 mM MgCl_2_, and 0.2 mg/ml BSA. Nsp1 and wHTH were added to the reactions as indicated in figure legends. The 16-mer DNA primer contains a Cy5 fluorophore attached to its 5’-end. The 73-mer template 5’-AATGATGAAGATATCT*GGTCGCTCC*ATTCT*GGAGCGACC*TCTTAATCTAAGCAC<u>TCGCTATGTTTTCAAG</u>TTT is prone to form the 9-bp hairpin (bases making the hairpin stem are shown in *italics*; the primer annealing site is underlined). Reactions were incubated for the indicated time points at 35 °C and stopped by mixing with equal volume of stopper solution containing 90% v/v formamide, 50 mM EDTA, pH 8, and 0.02% Bromophenol blue, heated at 95°C for 1 min, and resolved by 20% Urea-PAGE. The reaction products were visualized by imaging on Typhoon FLA 9500 (Cytiva) and quantified by ImageJ, version 1.54 (NIH). Experiments were repeated three times.

## CRediT authorship contribution statement

**Andrey G. Baranovskiy:** Conceptualization, Data curation, Formal analysis, Investigation, Methodology, Validation, Writing – review & editing, Writing – original draft. **Lucia M. Morstadt:** Investigation, Methodology. **Nigar D. Babayeva:** Investigation, Resources. **Tahir H. Tahirov:** Conceptualization, Funding acquisition, Project administration, Supervision, Writing – review & editing.

## Data availability

The results are available in Supplementary material.

## Acknowledgements

This work was supported by the National Institute of General Medical Sciences grant R35GM152032 to T.H.T. We thank J. Lovelace for assistance with wHTH/Polα model generation. AlphaFold-multimer modeling was performed at the Holland Computing Center of the University of Nebraska, which receives support from the UNL Office of Research and Economic Development, and the Nebraska Research Initiative.

