## Supplemental Figure 1 for "Nsp1 stalls DNA Polymerase α at DNA hairpins"

for the article

^1^Eppley Institute for Research in Cancer and Allied Diseases, Fred & Pamela Buffett Cancer Center. University of Nebraska Medical Center, Omaha, NE, USA.

**This file includes:**

Supplementary Figure S1

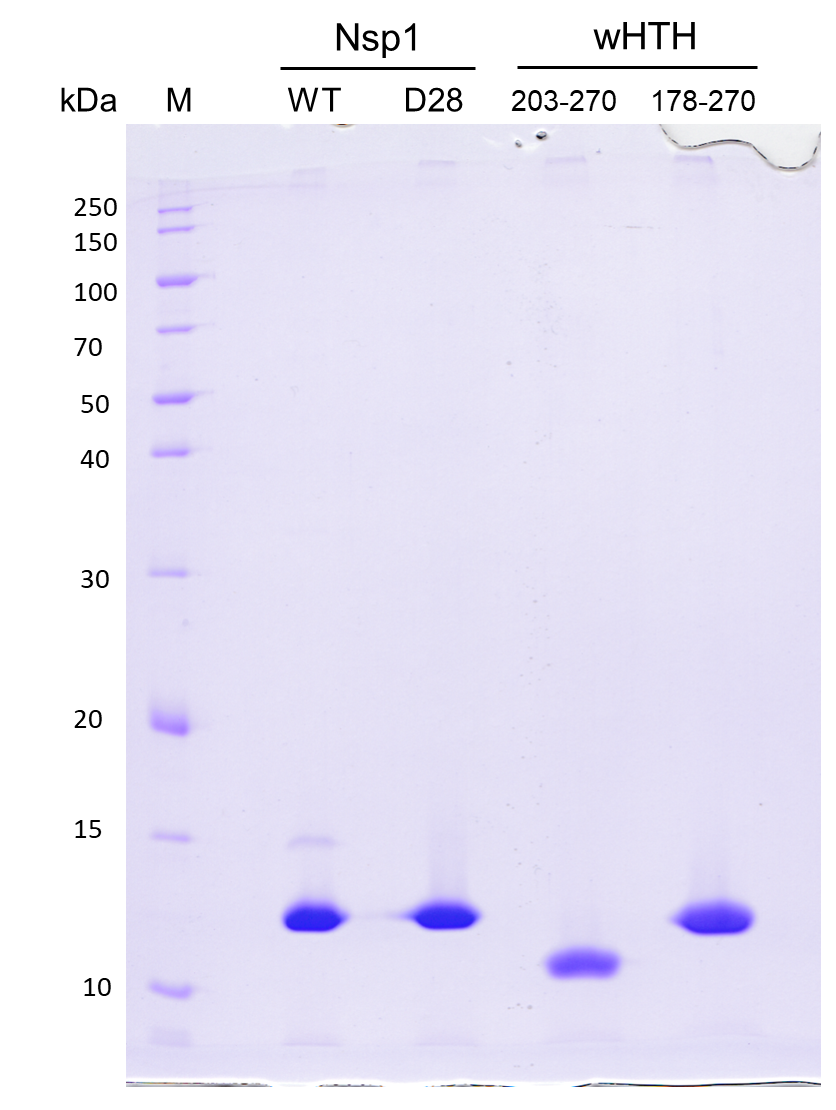

**Supplementary Figure S1. Analysis of purity of Nsp1, its mutant V28D, and two variants of wHTH.** The wHTH variant 178-270 was expressed and purified at significantly higher yield and chosen for functional studies. Proteins were separated by 13% SDS-PAGE and stained by Coomassie Brilliant Blue R-250.
